# SARM1 base-exchange inhibitors induce SARM1 activation and neurodegeneration at low doses

**DOI:** 10.1101/2025.03.25.644293

**Authors:** Anisha Mani, Mateusz Mendel, Paul Westwood, Claudia Bonardi, Wiebke Saal, Andreas Topp, Matthew Bilyard, Alessandro Brigo, Matthias Beat Wittwer, Jörg Benz, Bernd Kuhn, Achi Haider, Maude Giroud, James Keaney

## Abstract

SARM1 has emerged as a promising therapeutic target in neurology due to its central role in axonal degeneration and its amenability to different modes of small molecule inhibition. One chemical approach to modulate SARM1 involves orthosteric inhibition via a SARM1-mediated base-exchange reaction between a small molecule and nicotinamide adenine dinucleotide (NAD^+^), the substrate of SARM1, to generate the active inhibitor. Here, we report that subinhibitory concentrations of SARM1 base-exchange inhibitors (BEIs) paradoxically increase SARM1 activity and worsen SARM1-induced cell death and neuronal damage *in vitro.* Low dose administration of RO-7529, a SARM1 BEI, exacerbated experimental autoimmune encephalomyelitis (EAE)-induced neurodegeneration *in vivo*. Our data highlight a unique pharmacological feature of SARM1 BEIs that may limit their therapeutic application in disorders associated with SARM1 activation and axonal degeneration.

## Main Text

Axonal degeneration is a major clinical and pathological feature underlying neurodegeneration in multiple sclerosis, amyotrophic lateral sclerosis and other neurological conditions. SARM1, a nicotinamide adenine dinucleotide (NAD^+^) hydrolase, has emerged as a critical mediator of programmed axonal degeneration (1). Activation of SARM1 disrupts cellular energy homeostasis, depleting NAD^+^ and generating adenosine diphosphate ribose (ADPR), which perturbs intra-axonal calcium levels and induces axonal damage (2). SARM1 exists as a homomultimeric octamer with the autoinhibitory ARM domains locking the catalytic TIR domains in an inactive conformation (3). Both NAD^+^ and another metabolite nicotinamide mononucleotide (NMN) compete for binding to an allosteric site in the ARM domain (4, 5). During axonal injury it is posited that an increase in the NMN/NAD^+^ ratio and the higher affinity of NMN for SARM1 results in replacement of NAD^+^ in the allosteric pocket, releasing the ARM domains and allowing for TIR domain catalytic activity (6).

To date, multiple SARM1 inhibitors with diverse chemical structures have been reported including orthosteric (7, 8) and covalent allosteric (9) inhibitors. Orthosteric SARM1 inhibitors have been described which act via a base-exchange (BE) mechanism, in which the inhibitor (BEI) acts as a “pro-drug” and upon catalysis by SARM1, forms a covalent bond with ADPR to generate the active inhibitor (10). We previously discovered RO-7529, a potent SARM1 BEI (11). Intriguingly, we found that *in vivo* administration of RO-7529 at low doses exacerbated, and at high doses suppressed, plasma NfL levels in the spared nerve injury (SNI) model of acute peripheral nerve injury. To enhance our understanding of SARM1 BEI *in vivo* pharmacology, we assessed the activity of RO-7529 at a range of doses in the experimental autoimmune encephalomyelitis (EAE) model (Fig. 1A) - a chronic model of CNS autoimmune disease with associated axonal damage. Our aim was to select a low (2 mg/kg), medium (10 mg/kg) and high (50 mg/kg) dose that would cover a broad range of exposures covering sub-pharmacological to virtually complete target coverage (>IC_95_) (11). Starting at time of immunization, twice-daily oral administration of RO-7529 at 50 mg/kg reduced EAE symptoms, however RO-7529 at 2 mg/kg led to earlier EAE onset and worsened EAE during the acute inflammatory phase compared to vehicle-treated controls (Fig. 1B). Exacerbated EAE with low dose RO-7529 in the acute phase was accompanied by significant elevation in serum levels of neurofilament light chain (NfL), an established biomarker of axonal damage, compared to vehicle controls (Fig. 1C, D). In contrast, high dose RO-7529 suppressed serum NfL throughout the disease course (Fig. 1D, E). Histological assessments at study termination on day 28 indicated significantly increased inflammation and a trend for increased demyelination in the lumbar spinal cord of the low dose RO-7529 group (Fig. 1F-H). In addition to previous results in the SNI model (11), the EAE data represents a second disease model where we have observed worsening of axonal degeneration with a SARM1 BEI at low doses.

**Figure 1.**
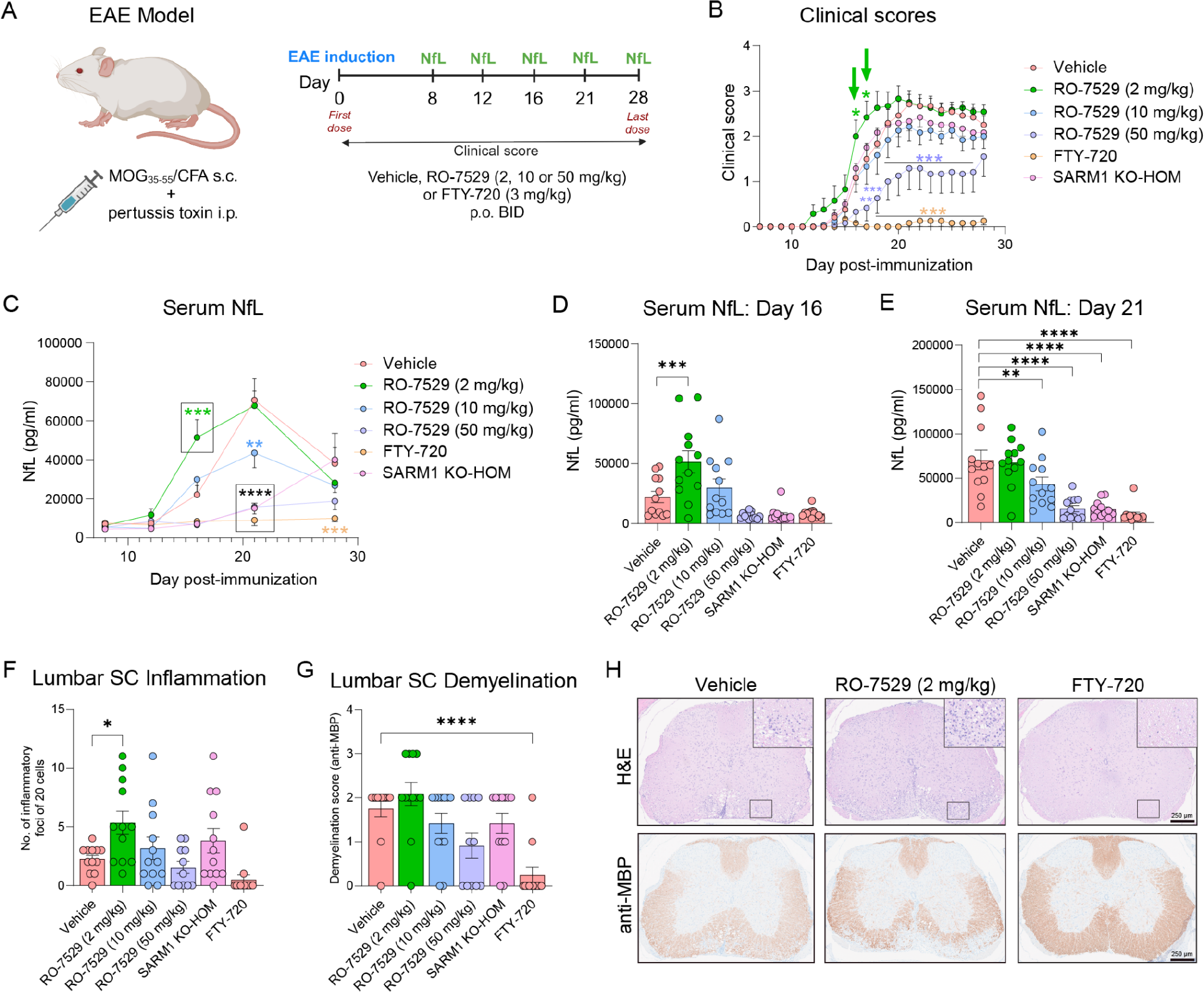
Low dose administration of a SARM1 BEI worsens neurodegeneration during the acute inflammatory phase of EAE. (A) Design of the EAE pharmacology study testing RO-7529 at 2, 10 and 50 mg/kg starting from time of immunization. (B) Mean disability clinical scores of EAE mice. Green arrows indicate worsening disability with RO-7529 at 2 mg/kg during the acute inflammatory phase. (C) Serum NfL levels in EAE mice on days 8, 12, 16, 21 and 28 post-immunization. Serum NfL levels at day 16 (D) and day 21 (E) from the same study. (F) Inflammation in lumbar spinal cord at day 28 quantified as the number of inflammatory foci containing at least 20 cells per section per mouse. (G) Demyelination in lumbar spinal cord at day 28 quantified using a demyelination scoring system based on anti-MBP staining. (H) Illustrative lumbar spinal cord sections stained with: *top* - hematoxylin (nuclei: blue) and eosin (cytoplasm: pink), *bottom* - anti-MBP (brown) and hematoxylin (nuclei: blue). Magnified white matter regions from H&E-stained sections demonstrate inflammatory infiltrates. The anti-MBP stained section from a RO-7529 (2 mg/kg)-treated mouse depicts a demyelination score of 3. For (B-G), n = 12 animals per group; data are shown as mean ± standard error of the mean (SEM). In (D-G), dots represent individual animals. Significance is indicated by ****P<0.0001, ***P < 0.001, **P< 0.01 and *P < 0.05, determined by two-way ANOVA and Dunnett’s post-hoc test.

To further understand the pharmacology of SARM1 BEIs and to explore the potential mechanism for low dose worsening of neurodegeneration, we next investigated diverse SARM1 BEIs (Fig. 2A) including previously published examples (7, 8) in biochemical and cellular assays of SARM1 activity, cell death and neurite degeneration. First, SARM1 NAD^+^ hydrolase (NADase) activity was measured in a cell-free biochemical assay via mass spectrometry detection of NAD and linear ADPR levels. At substrate concentrations similar to intracellular NAD levels (200 µM) (12), an increase in NADase activity was observed at subinhibitory concentrations of most SARM1 BEIs (Fig. 2B). Next, using human SY5Y neuroblastoma cells incubated with the neurotoxin vacor, a known SARM1 activator (13), we observed a significant increase in cell death at subinhibitory BEI concentrations but only in the presence of a subactivating vacor concentration of 5 µM (Fig. 2C). This effect was not observed in the absence of vacor or at vacor concentrations inducing almost complete cell death. Similarly, in human induced pluripotent stem cell (iPSC)-derived motor neurons incubated with vacor at 5 µM, low BEI concentrations significantly increased release of NfL relative to controls (Fig. 2D). In contrast, BEIs did not induce NfL elevation in the presence of vacor at 25 µM (Fig. 2E). NfL elevation upon incubation with both vacor at 5 µM and low BEI concentrations was also accompanied by observable neurite degeneration in iPSC-derived motor neuron cultures, as exemplified by Ex. 27 (Fig. 2F, G). In all three *in vitro* assays described, high BEI concentrations blocked SARM1 activity and reduced SARM1-mediated cell death and neuronal damage.

**Figure 2.**
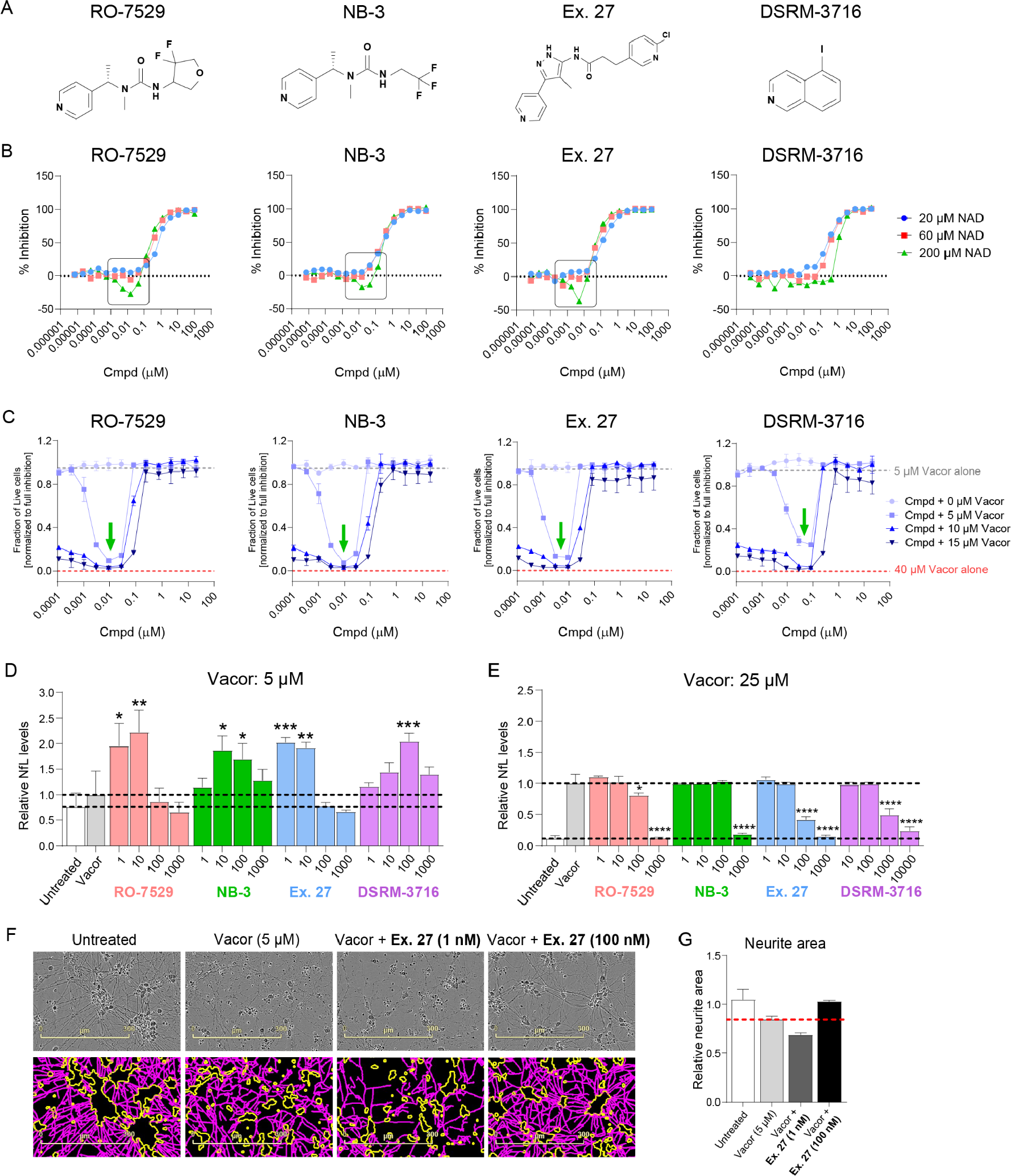
SARM1 BEIs induce SARM1 activation, cell death and neuronal damage at low doses *in vitro*. (A) Chemical structures of different SARM1 BEIs used in this study. (B) SARM1 NADase activity measured in a biochemical assay via mass spectrometry detection of NAD and linear ADPR levels. Substrate concentrations of 20, 60 and 200 µM NAD were assessed. The ratio of linear ADPR to NAD peak area was normalized against negative control (2% DMSO) and positive control (100 µM NB-3) and then plotted against compound concentration. Boxes indicate increased SARM1 activity. (C) SARM1 activation-induced cell death in SY5Y cells. Cell death following vacor exposure was assessed by measurement of ATP levels. Luminescent signals were normalized against untreated negative controls (0 µM vacor) and positive controls (40 µM vacor). Green arrows indicate increased cell death in the presence of both sub-µM BEI concentrations and a subactivating vacor concentration of 5 µM. NfL release from iPSC-derived motor neurons following exposure to 5 µM (D) or 25 µM (E) vacor and treatment with increasing concentrations of SARM1 BEIs. (F) Phase contrast (top) and Neurotrack (bottom) images of iPSC-derived motor neurons treated with vacor and low (1 nM) or high (100 nM) concentration of Ex. 27. Neurotrack images show the neurite mask in magenta and cell-body cluster mask in yellow. Images taken at 72 h post-vacor exposure. (G) Relative neurite area quantified using the Neurotrack module at 72 h post-vacor exposure. For (C-E) and (G), data are shown as mean ± standard deviation (SD) and results are representative of two to three independent experiments. Significance is indicated by ****P<0.0001, ***P < 0.001, **P< 0.01 and *P < 0.05, determined by one-way ANOVA and Dunnett’s post-hoc test.

To further delineate the role of base-exchange activity in SARM1-mediated neurotoxicity, we examined two mechanistically distinct small-molecule inhibitors, EV-99 and FK-866, both of which inhibit SARM1 through alternative pathways (Fig. S1A). EV-99 is an allosteric covalent inhibitor that binds to C311 in the ARM domain of SARM1, inhibiting enzymatic function without directly competing at the catalytic domain (9). FK-866, a potent nicotinamide phosphoribosyltransferase (NAMPT) inhibitor, blocks the conversion of NAM to NMN in the NAD^+^ salvage pathway, preventing increase in NMN levels and SARM1 activation (14). These inhibitors provide a valuable pharmacological contrast to SARM1 BEIs, allowing us to assess whether the toxicity observed at low BEI concentrations is a specific consequence of targeting the base-exchange mechanism. Crucially, neither EV-99 nor FK-866 exhibited any toxicity at low concentrations and a subactivating vacor concentration in human SY5Y neuroblastoma cells, conditions that led to cell death with subinhibitory BEI treatment (Fig. S1B). This lack of toxicity strongly suggests that the low dose exacerbation of neuronal damage is a specific effect driven by base-exchange activity.

Our findings may have implications for the development of SARM1 BEIs in treating neurological disorders associated with axonal degeneration. These *in vitro* and *in vivo* results indicate that while efficacious at high concentrations, SARM1 BEIs can induce SARM1 activation and neurodegeneration at subinhibitory concentrations and in conditions of partial SARM1 activation. In contrast, no BEI-induced cytotoxicity or neurite damage was observed in the absence of SARM1 activation in our cellular assays suggesting that safety evaluations in healthy animals or individuals may not identify this pharmacological effect. Measuring levels of SARM1 enzymatic activity during the course of human neurological disease will be crucial to determine the clinical risk of this low dose BEI effect. At the time of writing, our Genentech colleagues have reported similar findings of substoichiometric activation induced by SARM1 BEIs in mild SARM1-activating conditions *in vitro* and following acute dosing in acute neurotoxin and peripheral nerve injury models *in vivo* (15). While the mechanism for low dose activation remains to be fully elucidated, a new preprint suggests that ADPR adducts, including those generated from BEIs, can act as molecular glues to promote formation of superhelical SARM1 filaments which precipitate as condensates with full NADase activity (16). How this liquid-to-solid phase transition explains low dose SARM1 activation and high dose SARM1 inhibition with BEIs is currently unclear. In addition to the low dose effects, differences were observed between high dose RO-7529 and SARM1 knockout in the EAE model on both clinical score and serum NfL readouts (see Figs. 1B and 1C), which warrants additional investigation for potential off-target effects of RO-7529 and other SARM1 BEIs at high doses. Collectively, our findings highlight the challenges associated with SARM1 BEIs, emphasizing the need to avoid potential low dose paradoxical activation while informing human dosing regimens. Further, these results underscore the translational importance of incorporating biomarkers such as plasma NfL into clinical trial designs to mitigate risks and maximize the neuroprotective potential of SARM1 inhibitors in neurodegenerative diseases.

## Methods

### Compounds

Commercially available reagents were purchased at reagent grade and used without further purification. Dry solvents for reactions were reagent-grade, purchased from commercial suppliers, and used without further purification. LC-MS high resolution spectra were recorded with an Agilent LC-system consisting of Agilent 1290 high pressure gradient system, and an Agilent 6545 QTOF. The separation was achieved on a Zorbax Eclipse Plus C18, 1.7 µm, 2.1 x 50 mm column (P/N 959731-902) at 55 °C; A = 0.01% formic acid in Water; B= acetonitrile at flow 0.8 mL/min. gradient: 0 min 5% B, 0.3 min 5% B, 4.5 min 99 % B, 5 min 99% B. The injection volume was 2 µL. Ionization was performed in an Agilent Multimode source. The mass spectrometer was run in “2 GHz extended dynamic range” mode, resulting in a resolution of about 20 000 at *m*/*z* = 922. Mass accuracy was ensured by internal drift correction. LC–MS analyses to determine purities and associated mass ions were performed using an Agilent Infinity 1260 liquid chromatograph 6120 quadrupole mass spectrometer with positive and negative ion electrospray and ELS/UV at 254 nm detection using an Agilent Zorbax Extend C18, Rapid Resolution HT 1.8 μm C18 30 mm × 4.6 mm column, and a 2.5 mL/min flow rate with a 4 min run time. Nuclear magnetic resonance (NMR) spectra were recorded using a 500 MHz Bruker Avance NEO spectrometer or on a Bruker Avance III, 600 MHz spectrometer, both equipped with a 5 mm BBO (H&F) Z-gradient CryoProbe, in DMSO-*d*6 at 25 °C, or on a Varian Gemini 300, in DMSO-*d*6 or METHANOL-*d*4 at 25 °C. Chemical shifts were referenced to the residual solvent signals and are reported as *δ* [ppm], multiplicity, coupling constant *J* [Hz]. Multiplicities are denoted as s, singlet; d, doublet; t, triplet; q, quartet; quin, quintet; m, multiplet; br, broad, and combinations thereof.

***((3-[(3R)-4,4-difluorotetrahydrofuran-3-yl]-1-methyl-1-[(1S)-1-(4-pyridyl)ethyl]urea or 3-[(3S)-4,4-difluorotetrahydrofuran-3-yl]-1-methyl-1-[(1S)-1-(4-pyridyl)ethyl]urea)*** (Compound 7, RO-7529) was prepared as described in (11).

^1^H NMR (DMSO-*d*6, 500 MHz) *δ* 8.51 - 8.56 (m, 2H), 7.19 - 7.23 (m, 2H), 6.64 (d, 1H, *J* = 8.4 Hz), 5.51 (q, 1H, *J* = 7.1 Hz), 4.51 - 4.63 (m, 1H), 4.22 (dd, 1H, *J* = 8.8, 8.8 Hz), 4.02 - 4.11 (m, 1H), 3.78 - 3.88 (m, 1H), 3.72 - 3.78 (m, 1H), 2.60 (s, 3H), 1.43 ppm (d, 3H, *J* = 7.1 Hz); ^13^C NMR (DMSO-*d*6, 126 MHz) *δ* 157.5, 150.8, 149.7, 126.5 (dd, *J* = 255.4, 253.0 Hz), 121.8, 71.2 (dd, *J* = 30.9, 30.9 Hz), 70.0 (dd, *J* = 5.4, 1.8 Hz), 54.4 (dd, *J* = 30.9, 18.0 Hz), 50.9, 28.7, 15.9 ppm; ^19^F NMR (DMSO-*d*6, 471 MHz) *δ* –103.2 - (−)102.0 (m, 1F), –116.3 - (−)115.5 ppm (m, 1F); LC-HRMS (*m*/*z*): [M+H]^+^ calcd for C13H17F2N3O2 + H^+^: 286.1361, found: 286.1360, Diff −0.1 mDa.

***1-Methyl-1-[(1S)-1-(4-pyridyl)ethyl]-3-(2,2,2-trifluoroethyl)urea*** (NB-3) was prepared as described in (8). ^1^H NMR (300 MHz, METHANOL-*d*4) *δ* 8.59 - 8.42 (m, 2H), 7.51 - 7.22 (m, 2H), 5.64 (q, *J* = 7.1 Hz, 1H), 3.93 (q, *J* = 9.4 Hz, 2H), 2.70 (s, 3H), 1.58 ppm (d, *J* = 7.3 Hz, 3H); MS (ESI): m/z = 262.1 [M + H]^+^.

***3-(6-Chloro-3-pyridyl)-N-[4-methyl-3-(4-pyridyl)-1H-pyrazol-5-yl]propanamide*** (Ex. 27) was prepared as described in (17). ^1^H NMR (600 MHz, DMSO-*d*6) *δ* 13.15 - 12.83 (m, 1H), 10.01 - 9.66 (m, 1H), 8.63 (br d, *J* = 5.0 Hz, 2H), 8.33 (d, *J* = 2.3 Hz, 1H), 7.78 (dd, *J* = 2.5, 8.2 Hz, 1H), 7.59 (br d, *J* = 5.0 Hz, 2H), 7.45 (d, *J* = 8.2 Hz, 1H), 2.94 (t, *J* = 7.4 Hz, 2H), 2.68 (br t, *J* = 7.4 Hz, 2H), 1.96 ppm (br s, 3H); LC-HRMS (*m*/*z*): [M + H]^+^ calcd for C17H16ClN5O + H^+^: 342.1115, found: 342.1114, Diff −0.1 mDa.

EV-99 was prepared as described in (18). ^1^H NMR (300 MHz, DMSO-d 6) δ = 10.88 (s, 1H), 7.53 (d, J = 7.5 Hz, 1H), 7.29 (br d, J = 7.7 Hz, 1H), 7.18 - 6.74 (m, 6H), 6.69 (br s, 1H), 6.50 (br s, 1H), 6.26 (dd, J = 2.2, 16.7 Hz, 1H), 6.05 - 5.94 (m, 2H), 5.81 (br d, J = 10.3 Hz, 1H), 5.44 (br d, J = 6.6 Hz, 1H), 3.54 - 3.38 (m, 1H), 3.02 (br s, 4H); ESI-MS (ESI): m/z = 405.2 [M+H]+

### EAE model

#### Mice

Homozygous Sarm1-knockout (Sarm1-AGS3, hereafter SARM1 HOM-KO) and C57BL/6 female mice were used. SARM1 HOM-KO mice (JAX stock 034399) were bred at Charles River (Sulzfeld, Germany) for F. Hoffmann-La Roche Ltd., and C57BL/6 mice (JAX stock 000664) were bred and purchased from The Jackson Laboratory. Mice were 10-13 weeks old at disease induction and were group-housed in a controlled environment with regulated temperature, humidity, and a 12 h light/dark cycle, with unrestricted access to food and water. All the experiments were conducted under protocols approved by Hooke Laboratories’ IACUC, in compliance with the NIH Guide for the Care and Use of Laboratory Animals and PHS Policy on Humane Care and Use of Laboratory Animals.

#### EAE induction

EAE was induced in 60 female C57BL/6 mice and 12 female SARM1 HOM-KO mice, aged 10 to 13 weeks. Mice were acclimated for 7 to 9 days prior to the study. On Day 0, mice were immunized with MOG35-55/CFA emulsion (Hooke® Kit MOG35-55/CFA Emulsion PTX, catalog number EK-2110, Hooke Laboratories). The emulsion was injected subcutaneously at two sites: the upper back (1 cm caudal to the neckline) and the lower back (2 cm cranial to the base of the tail), with 0.1 mL per site. Pertussis toxin (140 ng in 0.1 mL) was administered intraperitoneally within 2 h of the emulsion injection and again 24 h later. EAE progression was monitored by weighing mice three times weekly (Monday, Wednesday, and Friday) starting from Day 0, and daily clinical scoring from Day 7 post-immunization by blinded individuals. The EAE scoring system ranged from 0 to 5: 0 - No motor function changes; 1 - Limp tail; 2 - Limp tail and hind leg weakness; 3 - Limp tail and complete hind leg paralysis, or limp tail with paralysis of one front and one hind leg, or severe head tilting, edge-walking, cage wall pushing, spinning when lifted; 4 - Limp tail, complete hind leg, and partial front leg paralysis (mice scoring 4 for two consecutive days were euthanized and given a score of 5 for the duration of the experiment); 5 - Complete hind and front leg paralysis with no movement, spontaneous rolling, or death due to paralysis. Intermediate scores were assigned when clinical signs did not precisely match the defined categories.

#### Compound administration

Twice-daily oral administration (10 mL/kg) of the test compounds began on Day 0 AM and continued through Day 28 AM, with doses given at the same time each day (+/− 1 h) and intervals of 10 to 14 h. Treatment groups included Vehicle (water) in C57BL/6 mice, RO-7529 at 2, 10, and 50 mg/kg in C57BL/6 mice, FTY-720 at 3 mg/kg (Fingolimod, a sphingosine-1-phosphate receptor (S1PR) modulator in 0.3 mg/mL ethanol; Cayman Chemical cat. no. 10006292) in C57BL/6 mice, and Vehicle (30% polyethylene glycol and 7% Soluplus® in water) in SARM1 HOM-KO mice.

#### Blood collection

Blood samples (∼50 µL) were collected from all mice at four timepoints (Days 8, 12, 16, and 21), with serum (∼25 µL) isolated and stored at −80°C for neurofilament light (NfL) analysis. Tail vein collection was used for the first timepoints, followed by retro-orbital sinus collection for subsequent timepoints. At study termination (Day 28), blood (∼100 µL) was collected via retro-orbital sinus under local anesthesia (proparacaine HCl ophthalmic solution 0.5%) administered prior to blood collection., and serum was stored at −80°C.

#### Serum neurofilament light chain (NfL) analysis

NfL analysis was performed on serum samples collected on Days 8, 12, 16, 21, and 28 using an assay protocol developed from (19). Serum samples were assayed in duplicate at one dilution (1:2). Wells were coated with UD1 (Uman Diagnostics, 27016, UD1 mAb 47:3) at 1.25 µg/mL (30 µL/well) and incubated overnight at 4°C. Wells were blocked with 1X TBS + 3% bovine serum albumin (BSA) (100 µL/well) and incubated overnight at 4°C. Samples and standard were incubated overnight at 4 C. Detection antibody UD3 (Uman Diagnostics, 27018, UD3 biotin-labelled mAb 2:1) was added at 0.5 µg/mL (25 µL/well) and incubated at RT for 1 h with shaking. SULFO-TAG labeled streptavidin (Meso Scale Discovery, R32AD) was added at 0.25 µg/mL (25 µL/well) and incubated at RT for 1 h while shaking. MSD GOLD Read Buffer B (Meso Scale Discovery, R60AM) was added at 150 µL/well, then read on the MSD instrument (Meso Scale Discovery, MSD QuickPlex SQ 120MM). Wash buffer was 1X TBS + 0.1% Tween 20. Assay diluent (for standards/samples, detection antibody, and streptavidin) was 1X TBS + 1% BSA + 0.1% Tween 20. NfL concentrations were calculated using a four-parameter logistic (4PL) fit with a standard made from reconstituted bovine NfL (Progen, 62208).

#### Tissue collection, processing and staining

Mice were euthanized using CO_2_ inhalation and transcardially perfused with PBS. Intact spinal cords were collected within the vertebral column and immersion fixed in 10% neutral buffered formalin for 48 to 72 h. After fixation, the vertebral columns were decalcified in Immunocal solution (Statlab, McKinney, TX) for 12 to 24 h before dissection into individual cervical (C3/C4), thoracic (T2/T4), and lumbar (L1/L2) segments. All 3 spinal cord segments from each animal were embedded together within their respective vertebral body sections and processed using standard methods to paraffin blocks. Slides were prepared using 4 µm paraffin sections (1 section/slide), each section containing C, T, and L spinal cord segments, and were stained with hematoxylin and eosin (H&E).

#### Immunohistochemistry (IHC) for myelin basic protein (MBP)

IHC for MBP was performed on a Ventana Discovery Ultra immunostainer. Slides were prepared using 4 µm paraffin sections (1 section/slide) and were deparaffinized online before heat-induced epitope retrieval (HIER) using Ventana Discovery CC1 HIER reagent for 8 min at 100 C followed by non-specific protein blocking. Sections were incubated with the primary antibody (rabbit anti-MBP clone EPR21188, Abcam, Cambridge, UK) for 32 min at 1:4000 dilution followed by 32 min with biotinylated anti-rabbit antibody at 1:1000 dilution. Detection of immuno-reactive end-product was performed with the Ventana Discovery DABmap kit, using the brown chromogen diaminobenzidine (DAB). Sections were counterstained with hematoxylin.

#### Histopathological evaluation

Histopathological analysis with subjective scoring for inflammation and demyelination was performed by a pathologist who was blind to group and clinical score. The severity of inflammation was scored by quantifying the number of inflammatory foci consisting of 20 or more immune cells on H&E-stained sections. Estimates of the number of inflammatory foci were made when the inflammatory infiltrates contained more than 20 immune cells. Subjective assessment for evidence of demyelination was performed using paraffin sections of spinal cord that had been labeled for MBP by IHC. The extent and severity of demyelination was based on an estimate of the total white matter (WM) area that had reduced DAB labeling intensity as follows: 0, no demyelination (less than 2% of WM with reduced DAB labeling); 1 (2 to 5% demyelinated area); 2 (5 to 20% demyelinated area); 3 (20 to 30% demyelinated area); 4 (30 to 50% demyelinated area); 5 (>50% demyelinated area).

### Cell culture

Human neuroblastoma SH-SY5Y cells (Roche RICB) were cultured in DMEM high-glucose medium (Gibco, 11965092) supplemented with 10% fetal bovine serum (FBS; Gibco, A3160502) and 1% penicillin-streptomycin (Gibco, 15140122) under standard conditions (37°C, 5% CO₂). Cells were split at 80–90% confluence at a ratio of 1:3 to 1:5. For the assay, cells were detached using 0.05% trypsin-EDTA (Gibco, 25300054) for 2–3 min at 37°C, resuspended in DMEM CM, and pelleted by centrifugation (300 × g, 5 min, room temperature). The cell pellet was washed with PBS, resuspended in Opti-MEM without phenol red (Gibco, 11058-021; phenol red interferes with the assay) and used for the cell death assay. Human iPSC-derived motor neurons (iCell motor neurons) along with their media and supplements (iCell Motor Neurons kit, 01279, Catalog R1049) were purchased from Fujifilm Cellular Dynamics, Inc. (FCDI, Madison, WI, USA). Cells were cultured according to the manufacturer’s instructions. The cells were seeded at a density of 10,000 cells per well in a poly-D-lysine pre-coated cell culture 384-well plate (Corning® BioCoat® Poly-D-Lysine 384-well Black/Clear Flat Bottom, 356663), which was then coated with laminin (Sigma-Aldrich, L2020-1MG) for 3 h at 37℃ at a concentration of 3 µg/mL in Dulbecco’s phosphate-buffered saline (DPBS) (Gibco, 14190-094). The cells were maintained in complete maintenance media (prepared by adding the kit’s supplements to the basal media) supplemented with 5 µM DAPT (Sigma-Aldrich, D5942-5MG, 20 mM stock prepared in DMSO) for the first week in culture, after which they were maintained only in complete maintenance media until Day 10. 50% of the culture medium was replaced with fresh medium every 2 days. Cell cultures were maintained at 37°C with 5% CO_2_.

### SARM1 NADase activity

Reactions were performed in a 10 µL volume consisting of 2.4 nM human SARM1 (28-724) (Abcam, ab271737), 20, 60 or 200 µM nicotinamide adenine dinculeotide (NAD) and 60 µM nicotinamide mononucleotide (NMN). All reagents were prepared in 25 mM HEPES pH 7.2, 50 mM NaCl, 1 mM EDTA and 0.0025% Tween20. To determine compound effect on the catalytic activity of SARM1 reactions were incubated for 40, 70 or 210 min at room temperature (20, 60 or 200 µM NAD, respectively) in the presence of a 16-point concentration range of compound (100 µM to 7 pM; 1 in 3 dilution between each point; 2% DMSO). Reactions were then quenched with 80 µL of 0.125% formic acid. Following reaction quenching the peak area of NAD and linear ADP ribose (ADPR) were measured by a RapidFire High Throughput Mass Spectrometry System (Agilent Technologies, Santa Clara, CA) using an API6500 triple quadrupole mass spectrometer (AB Sciex Framingham, MA). The ratio of linear ADPR to total NAD plus ADPR peak area was normalized against negative control (2% DMSO) and positive control (100 µM NB-3) and then plotted against compound concentration.

### SY5Y cell death assay

The assay was performed in a total reaction volume of 25 µL per well in 384-well solid white bottom polystyrene TC-treated microplates (Corning, 3570). Test compounds were prepared with a starting concentration of 20 µM and serially diluted 12 times in a 3-fold dilution series. Compound and control solutions were dispensed into assay plates (100 nL per well) using an Echo liquid handler. Following the addition of test compounds, SH-SY5Y cells (20,000 cells/well, 20 µL/well) were added to the plates. Plates were centrifuged at 100 × g for 1 min and incubated at 37°C for 60 min to allow compound pre-incubation. Vacor solution (5 µL/well), a SARM1 activator, was then added to achieve final concentrations of 0, 5, 10 and 15 µM, and plates were incubated for an additional 24 h at 37°C. At the end of the incubation period, 25 µL of CellTiter-Glo® 2.0 Reagent (Promega, G9242) was added to each well to lyse the cells and measure ATP levels. Plates were incubated for 10 min at room temperature in the dark to stabilize the luminescent signal. Luminescence, proportional to ATP content as an indicator of cell viability, was measured using a PHERAstar FSX Microplate Reader (BMG Labtech). Luminescent signals were normalized against untreated negative controls (0 µM vacor) and positive controls (40 µM vacor).

### Neurite degeneration assays in iPSC-derived motor neurons

On day 10 iCell motor neurons were treated with SARM1 inhibitors in a 4-point dose-response (1 to 1000 nM or 10 to 10,000 nM). Cells were pretreated with the compounds for 1 h at 37℃, followed by addition of the SARM1 activator vacor at a concentration of 5 or 25 µM for 72 h at 37℃. After 72 h of treatment, cell culture supernatants were collected and neurofilament light chain (NfL) levels in supernatants were measured using the NfL ELISA kit from Uman Diagnostics (20-8002). The NfL levels were normalized to that of the respective vacor control. The data analysis was performed using GraphPad Prism (version 10.3.0). The IncuCyte S3 live cell imaging system (Sartorius Göttingen, Germany) was used to image iCell motor neuron cultures. The images were analyzed using the IncuCyte Neurotrack module to detect neurites and cell bodies.

### Statistical analysis

Data and statistical analyses were performed using GraphPad Prism 9.0 software. Results are presented as means ± standard deviation unless otherwise stated. Data were analyzed using two-tailed Student’s t-test to compare between two groups. One-way or two-way ANOVA was performed, followed by Dunnett’s post hoc test for multiple comparison testing.

## List of reagents

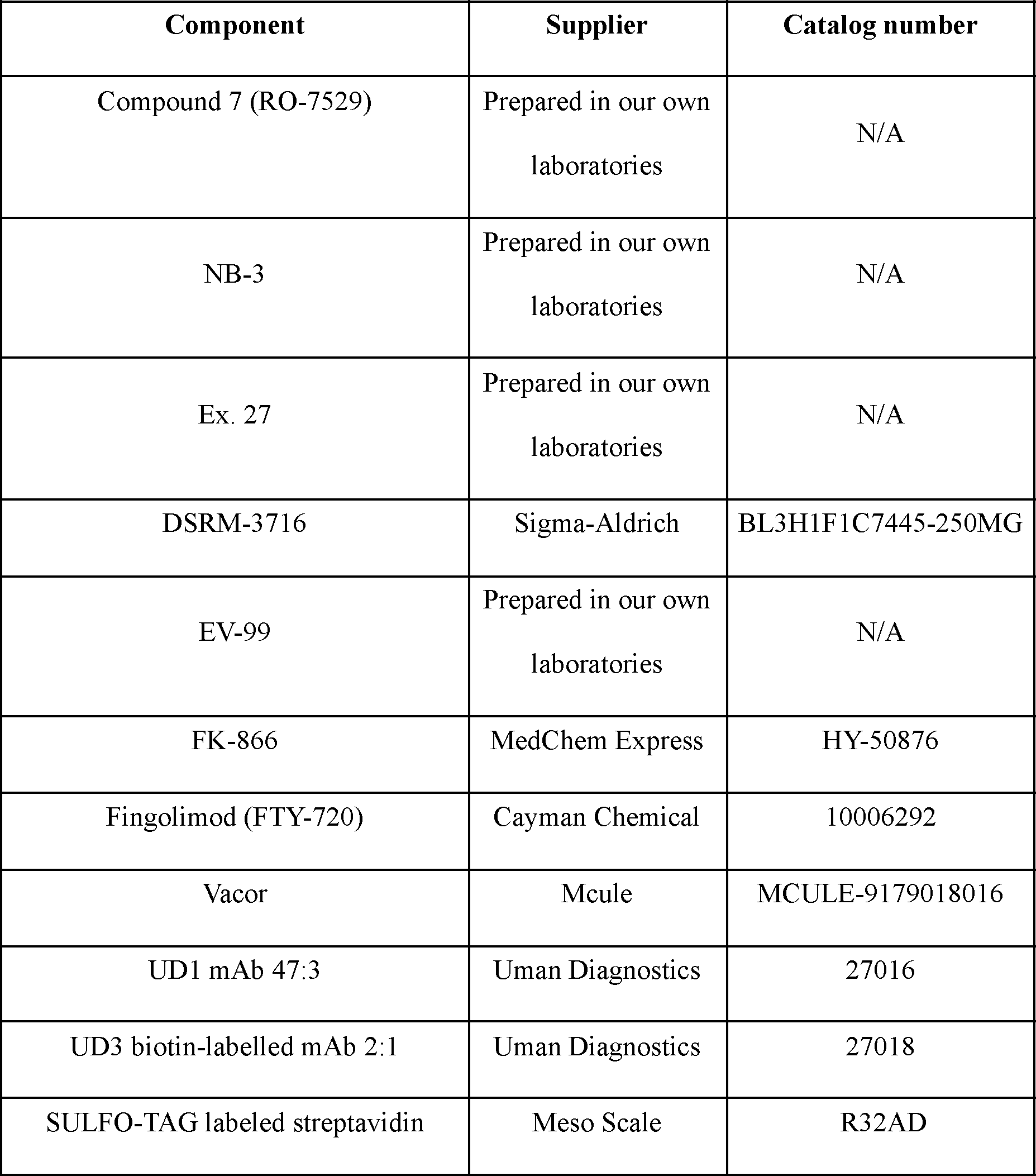
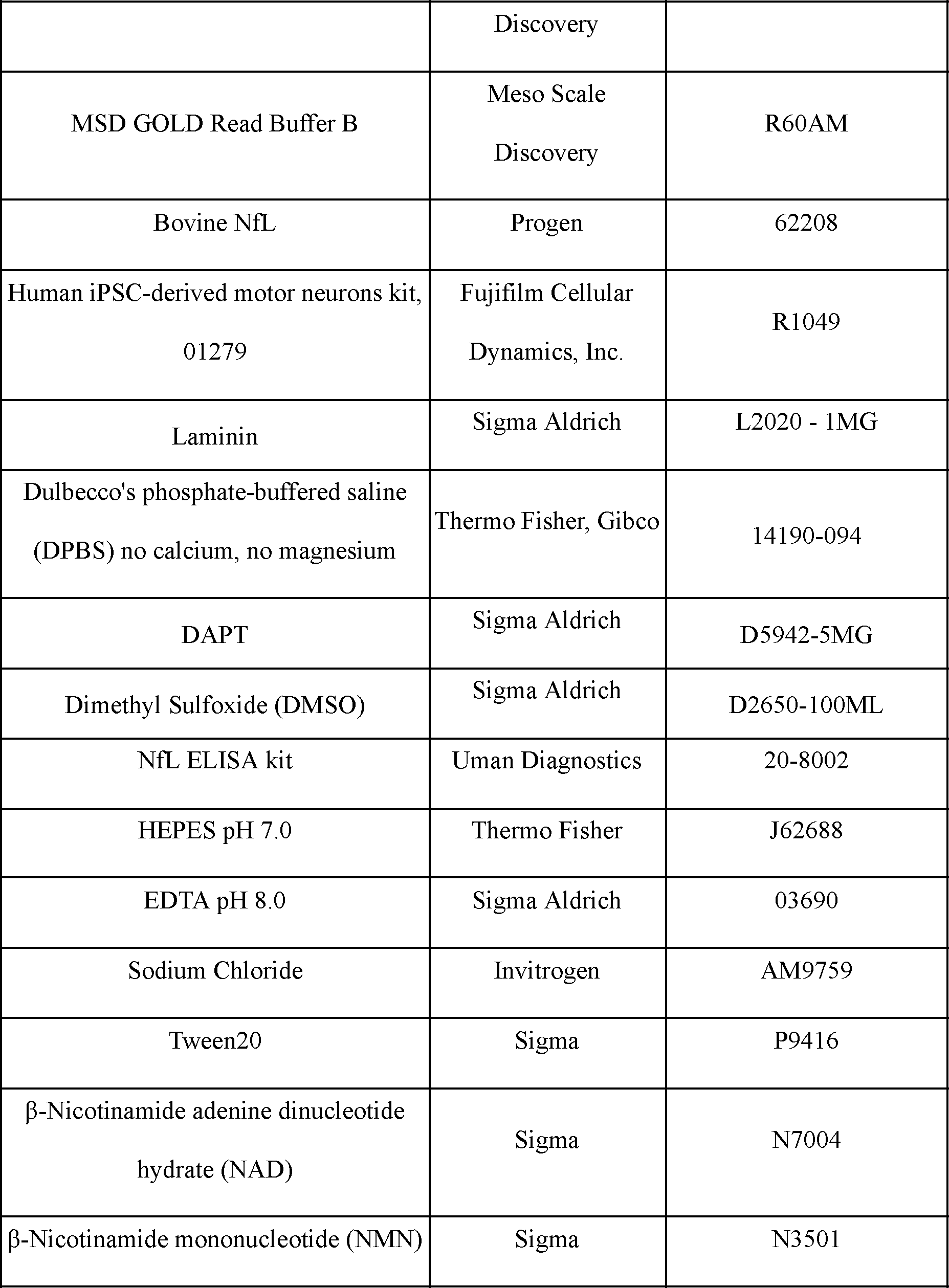
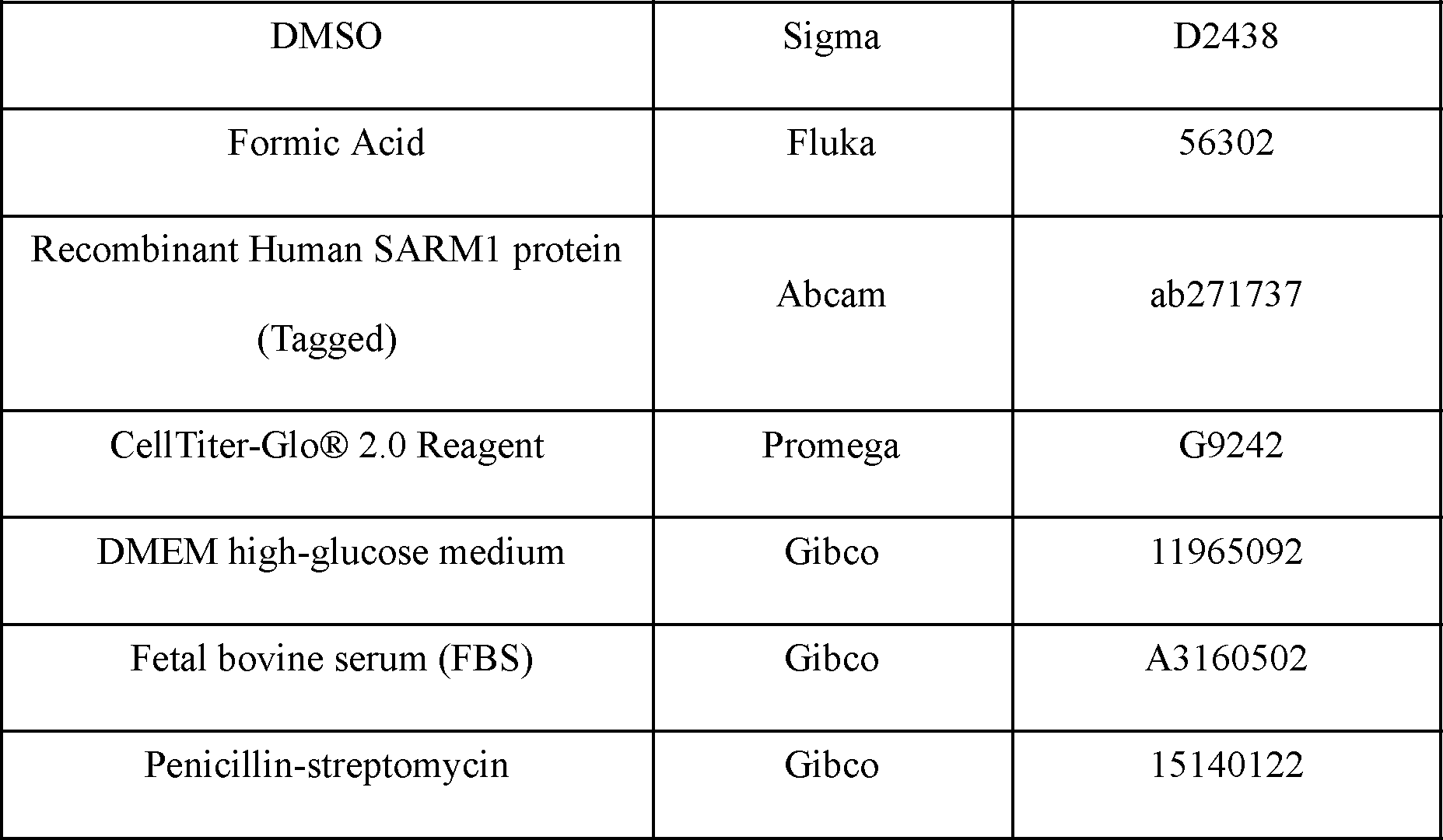

## Data Availability

All data generated or analysed during this study are included in this published article.

## Code Availability

Not applicable.

## Acknowledgements

All studies were funded by F. Hoffmann-La Roche Ltd, which supported the conceptualization, design, data collection, analysis and publication of this work. We thank Hooke Laboratories for conducting the mouse EAE study. Figure 1A was assembled using BioRender.

## Author contributions

AB, JB, BK, AH, MG and JK conceptualized the study. AM, MM, PW, AT and MB conducted the investigation. AM, MM, PW, CB, WS, AB, MBW, JB, BK, AH, MG and JK developed the methodology. Formal analysis was conducted by AM, MM, PW, CB, AB, MBW, JB, BK, AH, MG and JK. MG and JK had supervisory roles. The original manuscript draft was written by AM, MM and JK, and all authors read, revised and approved the manuscript.

