## Supplementary Figure 1 for "SARM1 base-exchange inhibitors induce SARM1 activation and neurodegeneration at low doses"

**This file contains: Supplementary Figure 1**

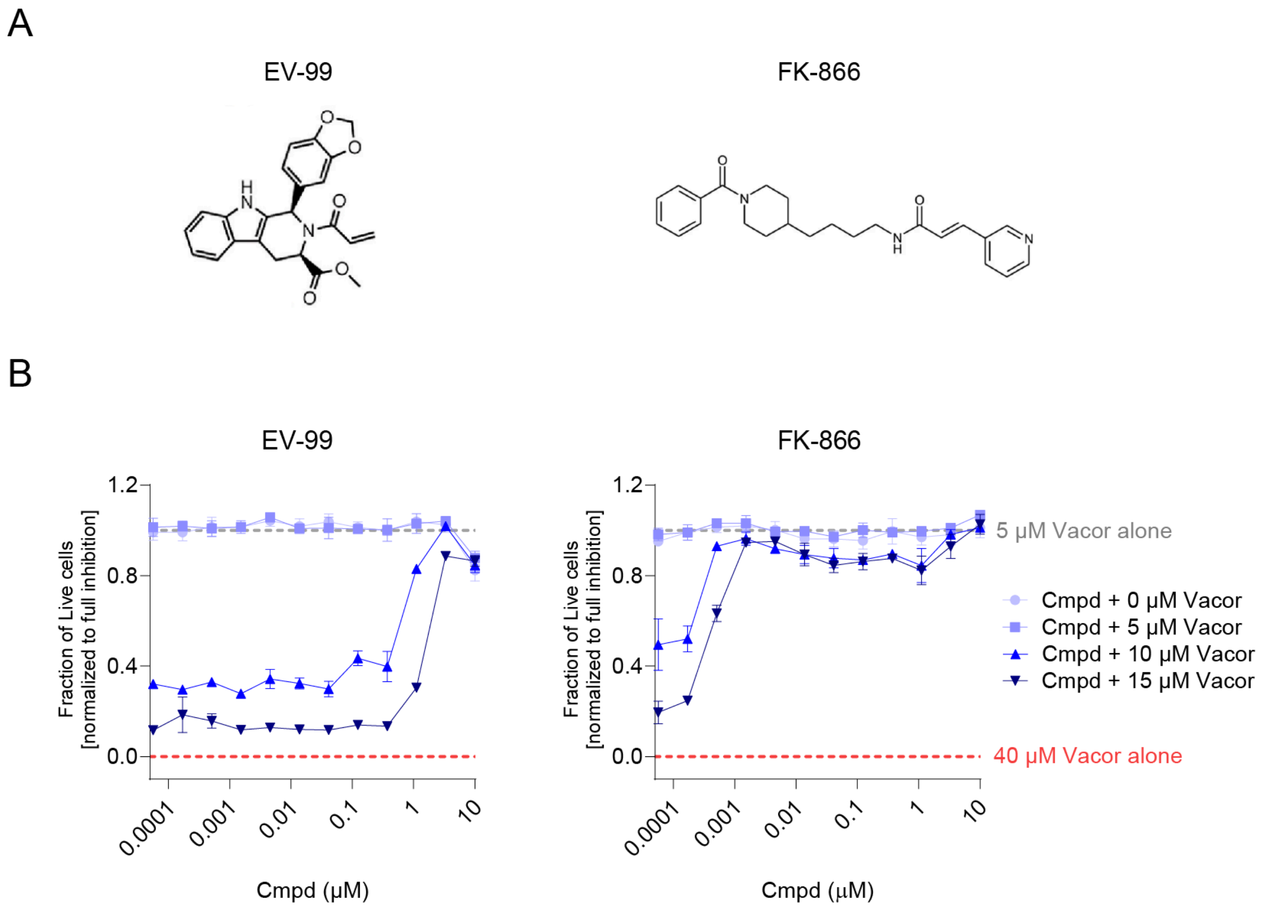


**Figure S1.** Non-orthosteric SARM1 inhibitors do not induce cell death at low doses *in vitro*. (A) Chemical structures of SARM1 allosteric covalent inhibitor (EV-99) and NAMPT inhibitor (FK-866). (B) Cell death assay in SY5Y cells shows no SARM1 activation-induced cell death at low inhibitor concentrations. Cell death following vacor exposure was assessed by measurement of ATP levels. Luminescent signals were normalized against untreated negative controls (0 µM vacor) and positive controls (40 µM vacor). Data is shown as mean ± standard deviation (SD) and results are representative of two independent experiments.
